# GrigoraSNPs: Optimized HTS DNA Forensic SNP Analysis

**DOI:** 10.1101/173716

**Authors:** Darrell O. Ricke, Anna Shcherbina, Adam Michaleas, Philip Fremont-Smith

## Abstract

High throughput DNA sequencing technologies enable improved characterization of forensic DNA samples enabling greater insights into DNA contributor(s). Current DNA forensics techniques rely upon allele sizing of short tandem repeats by capillary electrophoresis. High throughput sequencing enables forensic sample characterizations for large numbers of single nucleotide polymorphism loci. The slowest computational component of the DNA forensics analysis pipeline is the characterization of raw sequence data. This paper optimizes the SNP calling module of the DNA analysis pipeline with runtime results that scale linearly with the number of HTS sequences (patent pending)[1]. GrigoraSNPs can analyze 100 million reads in less than 5 minutes using 3 threads on a 4.0 GHz Intel i7-6700K laptop CPU.

## I. INTRODUCTION

Rapid analysis of DNA forensics samples can aid investigations. The application of high throughput sequencing (HTS) to DNA forensics samples is providing additional forensics capabilities, including: improved kinship detection, mixture analysis, biogeographic ancestry prediction, phenotype/externally visible traits (EVTs) prediction, analysis of trace DNA samples, and more. HTS DNA sequencing enables the sequencing of short tandem repeats (STRs) and single nucleotide polymorphisms (SNPs). Sizing of STR alleles by capillary electrophoresis is the current standard for criminal DNA forensics methods[2]. HTS enables sequencing loci alleles[3], expansion of panels to additional loci, and improved detection of contributor samples with lower DNA concentrations. Samples can be multiplexed for increased throughput and decreased cost per sample by labeling of samples with DNA barcodes. Reducing the time to analyze HTS DNA samples will aid forensic investigations.

DNA forensics is starting to shift to larger and sometimes hybrid STR and SNP panels. The FBI CODIS[4] system just expanded from 13 STR loci to 20 to include additional loci used in Europe. Rapid STR analysis[5] is starting to aid forensic investigators. The Illumina ForenSeq HTS panel consists of 68 STRs and 172 SNPs[6]. In addition to all CODIS loci, ForenSeq includes additional X and Y chromosome STRs, 94 SNPs for identity, 56 SNPs for biogeographic ancestry (BGA) prediction, and 22 SNPs for phenotype prediction[6]. Larger panels of loci have been developed with SNPs targeted for the analysis of complex DNA mixtures, kinship, externally visible traits (EVT)/phenotype, and more[7–9]. Even larger panels of SNPs are possible with DNA microarrays. Measurements of millions of SNPs are possible with DNA microarrays[10, 11]. These microarrays currently require more DNA than HTS and cannot yet be used for trace DNA forensics samples. However, the characterization of millions of SNPs could drastically improve kinship and biogeographic predictions. Like rapid STR analysis, rapid profiling and analysis of HTS and possibly microarray samples will impact forensic investigations. Rapid sequencing approaches like nanopore sequencing[12] appear very promising, but improvements in the current base calling error rates[13] are needed for forensic applications.

The bioinformatics community has developed a rich set of HTS data SNP analysis tools: SAMtools[14], GATK[15]^-^[16], SOAPsnp[17], SVNMix2[18], VarScan[19], MAQ[20], and more. Many of these tools were designed for variant analysis of exome and whole genome shotgun (WGS) sequences. Most of these tools rely upon the alignment of HTS sequences to a reference genome using very fast alignment tools (BWA[21] or Bowtie2[22]) based on the Burrows-Wheeler Aligner[23]. Standard pipelines based on SAMtools and GATK are very popular in the bioinformatics community. However, these tools are not optimized for the analysis of HTS SNP panels. For forensic HTS SNP analysis, the tool ExactID has been developed[ref]. While fast, ExactID relies upon primer sequences to assign reads to SNP loci. Lincoln Laboratory has developed multiple HTS SNP panels for kinship[7] and mixture analysis[9] using the Ion Torrent Proton and S5 platforms. The standard HTS sequences from these panels lack primer sequences. The HTS STR allele identification tool, STRait Razor, has been recently enhanced to tally SNP loci sequences as a step towards SNP allele calling[24]. To be able to rapidly characterize these HTS SNP forensic panels, GrigoraSNPs was developed. With 3 threads, GrigoraSNPs can analyze 100 million multiplexed HTS sequences using 3 threads on a 4.0GHz Intel i7-6700K laptop in less than 5 minutes. Technology advances with nanopore or other HTS technologies will enable real-time portable DNA forensics.

## II. METHODS

### A. SNP Panels

SNP panels are designed with multiple pairs of oligonucleotide primers that each amplifies target location(s) in the genome for characterization of SNP alleles. Each HTS sequence usually contain the following components: multiplexing barcodes on the 5’ and sometimes 3’ end of each sequence, 5’ and 3’ primers, and the SNP surrounded by flanking DNA sequences (Figure 1). HTS DNA datasets also contain other sequences arising from sequencing and polymerase chain reaction (PCR) amplification artifacts. For Thermal Fisher Scientific Proton and S5 HTS SNP sequences, the 5’ barcode predominantly start at the first or second base pair positions in sequences. Also, the 5’ and 3’ primer sequences are not present in the sequences generated. Multiple HTS SNP panels have been designed by MIT Lincoln Laboratory and evaluated with the GrigoraSNPs analysis tool for SNP allele calling.

**Figure 1.**
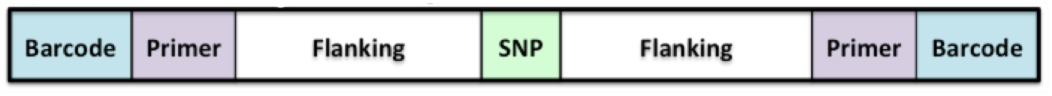
HTS DNA Sequence Components

### B. GrigoraSNPs

GrigoraSNPs was developed in Scala leveraging the Akka Actors model for efficient parallel processing on symmetric multiprocessing (SMP) computers. With a SNP dataset of N million sequences of length L for a panel of M amplification loci has the computational complexity of O(L x N x M). Popular tools replace the M targets with the entire human genome, further increasing the complexity of the problem. The standard multiplexed Ion Torrent SNP panel HTS sequences lack primer sequences for rapid SNP loci identification. To enable rapid loci identification, GrigoraSNPs was designed with the addition of a loci lookup table (tags) of 12 base pairs selected immediately following the 5’ barcode and adaptor sequence (linked bases GAT). This reduces the complexity of panel size from O(M) to O(1). Proton/S5 platform optimal results are obtained with 4 tags (2 per strand) for each target loci because of two different dominant 5’ ends for each target loci for each strand orientation. A utility tool, FindTags was developed in Scala for identifying tags from experimental data. A panel of 14,938 SNP loci was used for benchmarks. A Tag file with 87,531 tags was generated for the benchmark panel. Loading this tag file into memory requires 2.2 megabytes (MB) of memory. GrigoraSNPs uses reference target sequences from dbSNP[25] and optimizes the loci identification problem with a lookup table to ignore the size of the target SNP panel. By chance, some loci can share tag sequences. Some tags map to multiple loci, each candidate target is compared to HTS reads with matching tags. Each sequence is compared against the specified set of barcodes used for the experiment (sequencing run). If the entire barcode file is selected, GrigoraSNPs will automatically identify which barcodes were used (auto barcode detection – default). After a sequence matches a tag, specific SNPs are identified by matching the 10 nucleotides immediately flanking each SNP to the reference target sequences from dbSNP. To avoid incorrect SNP identification, GrigoraSNPs requires a minimum match of 19 or 20 bases flanking the target SNP. A very small error rate for SNP allele identification was observed with a minimum match threshold of 18 of 20 bases (data not shown).

Performance testing was done on two systems: HP DL380 with 2 CPUs and a SAGER laptop. The HP DL380 has two 2.3GHz Xeon 2698 v3 CPUs with 16 cores each (64 threads total), 512 GB RAM, 10 TB RAID using 10K RPM disks. The SAGER laptop is configured with Intel i7-6700K 4.0GHz CPUs with 4 cores (8 threads), 64 GB RAM memory, and a mirrored RAID pair of 1 TB Samsung 850 EVO SSD.

### C. SAMtools Pipeline

The dominant SNP analysis tool in the bioinformatics community is SAMtools. To characterize the results of GrigoraSNPs, a SAMtools pipeline was developed for SNP analysis. BWA[21] Mem(v0.7.12) was used to map each subject. SAMtools[14] fixmate (v1.3) was used to convert SAM output file to BAM. SAMtools sort (v1.3) was used to sort the BAM files into coordinate order. SAMtools Index (v1.3) was used to create an index file for the sorted BAM file. To produce variant calls using SAMtools (v1.3), a position file was created that contained the chromosome and position of each SNP of interest. SAMtools mpileup (v1.3) used this input file to create a BCF (binary variant call format) file of variants. SAMtools calls (v1.3) was used to parse and convert the file from BCF to VCF format. A parser was built for the VCF file with mapped rsids (reference SNP identifiers) and calculated total read depth, allele calls, and strand counts for each locus.

### D. GATK Pipeline

GATK is a popular sequence analysis tool developed by the Broad Institute. A GATK pipeline was also implemented for comparison for results with GrigoraSNPs and SAMtools. A GATK pipeline was developed for SNP analysis. BWA Mem was used to map each subject. SAMtools fixmate (v1.3) was used to convert SAM output file to BAM format. SAMtools[14] sort(v1.3) was used to sort the BAM files into coordinate order. SAMtools index (v1.3) was used to create an index file for the sorted BAM file. To produce variant calls using GATK[15, 16] Haplotype Caller (v3.5), BCFtools (4.2)[15] was used to create a VCF file from a list of loci rsids. Loci that did not have an rsid were manually added to the input VCF file. GATK HaplotypeCaller called variants only for loci in the input VCF file. A parser was built for the VCF file that determined total read depth, allele calls, and strand counts for each locus.

### E. STRait Razor

STRait Razor[24] is a popular STR HTS analysis tool. Version 3.0 includes identification of sequences containing SNP alleles. A custom STRait Razor configuration file was created based on dbSNP reference sequences for 14,833 of the 14.9k target SNPs with some Y chromosome loci not included due to small reference sequences (20 base pairs).

## III. RESULTS

HTS sequences from a HTS SNP panel (14.9k SNPs) were used to evaluate the performance of the SNP analysis pipelines. Figure 2 shows the performance results for 25, 50, 100, 250, 500, and 1,000 million sequences. The detected SNP counts for the SAMtools, GATK, and GrigoraSNPs tools are shown in Figures 3. The runtime for GATK using the –nct option (number of CPU threads to allocate per data thread) for 25M reads was 646 minutes using 3 threads compared to 290 minutes without this option; thus, all GATK runtimes shown are without the –nct option.

**Figure 2.**
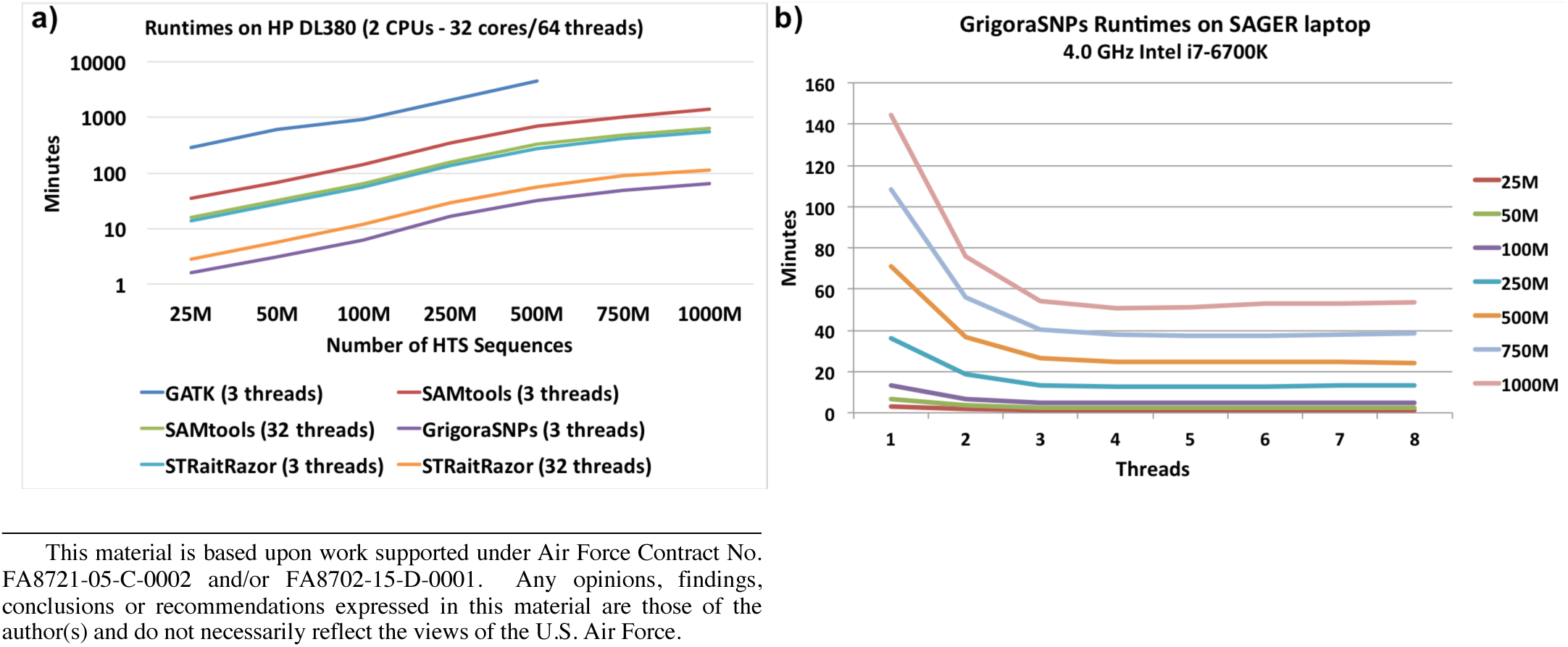
HTS SNP Analysis Performance Results. a) runtimes on HP DL380 with 3 threads and 32 threads for SAMtools; b) GrigoraSNPs runtimes on SAGER laptop with different numbers of threads.

**Figure 3.**
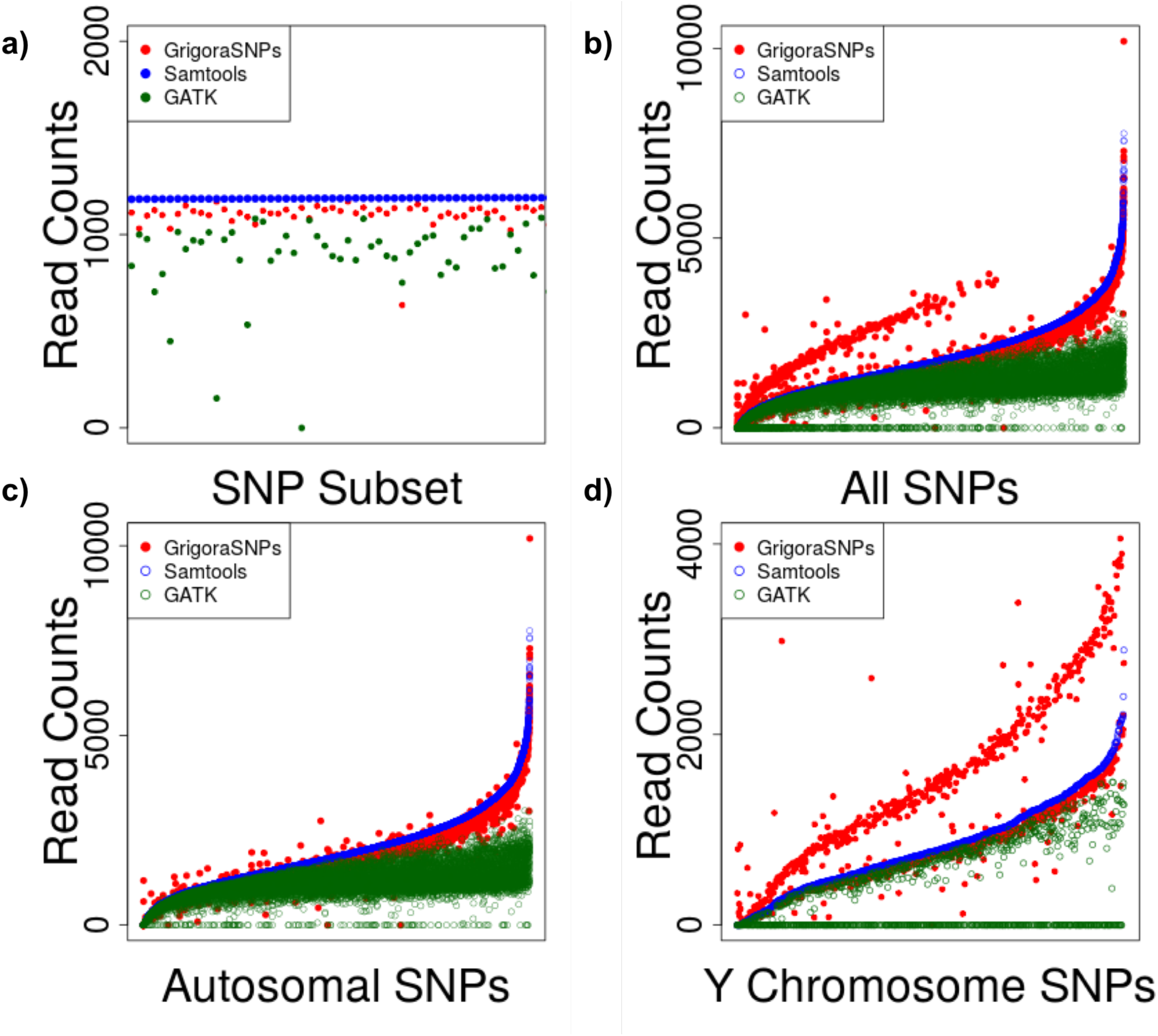
SNP counts for GrigoraSNPs, SAMtools, and GATK sorted by SAMtools counts. a) example 50 SNPs; b) all 14.9k SNPs; c) autosomal SNPs; and d) Y-chromosome SNPs.

## IV. DISCUSSION

The performance of GrigoraSNPs scales linearly with the number of sequences irrespective of SNP panel size (Figure 2). The lookup tag approach reduces the complexity of panel size from O(M) to O(1). The runtimes for GrigoraSNPs are very fast in comparison to GATK and SAMtools (Figure 2). The runtimes for GrigoraSNPs are 10 times faster than STRait Razor, more than 20 times faster than SAMtools, and 140 times faster than GATK with 3 threads (Figure 2). GrigoraSNPs processes 30 MB/sec HTS data, which is 3 times faster than ExactID[26]. Note that STRait Razor does not currently handle multiplexed HTS data or actually call SNP alleles. GrigoraSNPs reduces the runtime and computer size required to process forensics HTS DNA sequences. GrigoraSNPs enables DNA forensics on a laptop (Figure 2-b). GrigoraSNPs also reduces the free disk space requirements because it can characterize multiplexed sequence data without splitting the data into separate barcode files.

GrigoraSNPs characterizes roughly the same number of sequences per locus as SAMtools (Figure 3b). Random sequencing errors in some target HTS tag sequences likely cause the small loss of reads seen in Figure 3 for GrigoraSNPs when compared to SAMtools results. The expected loss is the instrument bse calling error rate times the length of the tag sequence times the number of HTS sequences for that loci. For example, an error rate of 0.1% times 10 bases would result in an expected average loss of 100 (i.e., 1%) of 10,0000 sequences. An individual with a rare sequence variant within a target tag sequence, would be expected to miss roughly 25% of the HTS sequences because the other 3 tag sequences for that loci will not be impacted. SNP call totals are shown in Figure 3 for GrigoraSNPs, SAMtools, and GATK pipelines. Note that GATK discontinued support for the Proton platform[27]. The SAMtools pipeline performs very well for all autosomal SNPs, but the BWA alignment to reference human genome step ends up mapping roughly 50% of the Y chromosome SNPs to the X chromosome in the pseudoautosomal region (PAR) (Figure 3). Note the large upper arc band of GrigoraSNPs SNP alleles compared to the sorted SAMtools allele calls in Figure 3b and 3d. These allele calls are approximately 2X the corresponding SAMtool values (Figure 3b – all SNPs and Figure 3d – Y chromosome SNPs) but are missing from only autosomal SNPs in Figure 3c.

GrigoraSNPs processes 100 million HTS reads in less than 5 minutes on an Intel 4 core laptop with 3 threads (Figure 2). Runtimes are not reduced on the DL380 by use of faster input and output with the HP workload accelerator; indicating that the application is not I/O bound. A 4.0 GHz Intel i7-6700K CPU system with 3 to 8 threads can run GrigoraSNPs in 5 minutes with the optimal performance seen for 3 threads at 4.7 minutes runtime (Figure 2).

HTS sequencing enables characterization of trace DNA samples, identification, improved kinship detection, mixture analysis, biogeographic ancestry prediction, externally visible traits prediction, and more. In DNA forensics, being able to go from sample to profile is fundamental. In some scenarios, being able to quickly characterize samples is essential for generating leads in a case. GrigoraSNPs eliminates a major time bottleneck for processing raw sequences to SNP allele calls.

## V. CONCLUSION

GrigoraSNPs provides an efficient solution for the rapid characterization of HTS DNA SNP sequences.

